# When to care and when to kill: termites shape their collective response based on stage of infection

**DOI:** 10.1101/287441

**Authors:** Hannah E. Davis, Stefania Meconcelli, Renate Radek, Dino P. McMahon

**Affiliations:** Institut für Biologie, Freie Universität Berlin, Königin-Luise-Str. 1-3, 14195 Berlin, Germany; Department for Materials and Environment, BAM Federal Institute for Materials Research and Testing, Unter den Eichen 87, 12205 Berlin, Germany

## Abstract

Termites defend their colonies from disease using an array of social behaviours, including allogrooming, cannibalism, and burial. We tested how groups of eastern subterranean termites (*Reticulitermes flavipes*) deploy these behaviours when presented with a nestmate at different stages of infection with the entomopathogenic fungus *Metarhizium anisopliae*. As expected, the termites groomed pathogen-exposed individuals significantly more than mock-treated controls; however, grooming levels were significantly higher after spore germination than before. Cannibalism became prevalent only after exposed termites became visibly ill, and burial was rarely observed. These results demonstrate that termites employ different strategies depending on the stage of infection that they encounter. Grooming intensity is linked not only to pathogen presence, but also to germination status, and, given the temporal correlation between cannibalism and visible signs of illness, the host may play a role in triggering its own sacrifice.

## Introduction

Social insects have evolved collective behaviours to protect their colonies from disease. These social immune defences, which include pathogen avoidance, prophylactic secretions, grooming, and corpse disposal, act to protect the colony as a whole, at times at the expense of individual members [1, 2]. In this latter case, sick colony members are identified and killed to prevent the spread of disease [1, 3, 4]. Regulation is therefore essential, both to prevent unnecessary killing and to allow the colony to dynamically adjust its investment in other defences [1].

Of all the social insects, the social Hymenoptera are the most well-studied. In honeybees (*Apis* spp. Linnaeus) activation of the physiological immune system by an infection results in a changed cuticular hydrocarbon profile [5], which can then trigger the removal of the infected bee by other members of the hive [6]. Likewise, workers respond to volatiles emitted by sick or injured brood by removing them from the hive [7, 8], and factors external to the host, such as the odour of a parasite or pathogen inside a brood cell [9], can also play a role. In ants, the situation is similar: invasive garden ant (*Lasius neglectus* Van Loon, Boomsma & Andrásfalvy) workers groom fungus-exposed pupae to prevent disease, but kill them if alerted to an internal infection by a change in cuticular hydrocarbons [4]. European fire ant (*Myrmica rubra* (Linnaeus)) workers also behave more aggressively toward fungus-infected adult nestmates once internal proliferation has begun [10].

Comparatively little is known about how termites (Blattodea: infraorder Isoptera) shape their social immune response based on the stage of infection encountered. There is broad consensus that the initial response to a pathogen-exposed nestmate is dominated by intense allogrooming [11–15], and cannibalism becomes more prevalent at some later stage [12, 16, 17]; however, when the switch occurs remains unclear. Although it has long been known that termites eat both the sick and the dead [18–20], with individuals most commonly eaten when “moribund but not yet dead” [3], no study to date has attempted to identify the stage of infection at which the risk of cannibalism first begins to increase. Should a termite die from an infection, or for any other reason, necromones attract worker termites to the corpse, which they preferentially eat (necrophagy) [21]. Corpses that are too old [22] or too numerous to consume [23] are defecated on and then buried, isolating them from the colony [18]. Burial of live individuals has also been observed [11, 24].

Each component of the social immune response serves to prevent a pathogen from reaching the next stage in its life cycle, and ultimately to prevent an epizootic, but it will only be effective if deployed at the appropriate time. In the specific case of *Metarhizium anisopliae* (Metchnikoff) Sorokin (Ascomycota: Hypocreales), a generalist entomopathogenic fungus, allogrooming is highly effective in removing most infectious conidia from the cuticle before they can germinate [12, 15, 25, 26]. Groomers can safely swallow the conidia [15, 27], and low-level infections acquired through contact with an infected nestmate may even boost individual anti-fungal defences [13]. Once an internal infection has been established, however, allogrooming is no longer effective [4]. The infected termite cannot be saved, and the longer it is left alive, the higher the colony-level fitness cost: resources that it consumes will go to support fungal growth, it will become increasingly unable to work, and should the fungus sporulate from its corpse, it will put the entire colony at risk [28].

We would therefore expect to see a switch from a grooming-dominated collective immune response to a cannibalism- and/or burial-dominated response beginning at the earliest point at which termites can detect a terminal infection. To address this, we used the eastern subterranean termite, *Reticulitermes flavipes* (Kollar), and the entomopathogenic fungus *M. anisopliae* to examine how the stage of infection encountered by a colony determines the collective response. Our hypotheses were that i) allogrooming would be most intense before conidial germination; ii) the shift to cannibalism would begin shortly after conidial germination. Contrary to expectations, we found that levels of grooming rose significantly after conidial germination, and that cannibalistic behaviours coincided with termite sickness, with a more rapid switch to cannibalism at later stages. By dividing the infection into stages [29] and studying how the social immune response differs over time, our study sheds new light on the processes by which social Blattodea identify fatally ill colony members and thereby defend their colonies from disease.

## Materials and Methods

### Insects

Three captive eastern subterranean termite (*Reticulitermes flavipes*) colonies at the Federal Institute for Materials Research and Testing (Bundesanstalt für Materialforschung und - prüfung, BAM) in Berlin, Germany were used in these experiments: colonies E, 5, and 8. Colony E was collected in Soulac-sur-Mer, France, in 2015. It was maintained in a dark room at 28°C, 83% humidity. Colonies 5 and 8 were collected in the vicinity of Le Grand-Village-Plage, Île d’Oléron, France, in 1994 and maintained in a separate dark room at 26°C, 84% humidity. Colonies were housed in physically separate sheet metal tanks as described by Günther Becker [30]. All three colonies appeared healthy.

Cardboard bait was used as the primary method for extracting termites from their parent colonies. From collection until staining or transfer to Petri dish nests, termites from the same colony were maintained in a plastic box containing cellulose pads (Pall Corporation, Port Washington, USA) moistened with tap water. Each colony box was maintained under the same temperature and humidity conditions as the parent colony, and tap water and new cellulose pads were added as needed.

### Fungi

The entomopathogenic fungus *Metarhizium anisopliae* DSM 1490 was maintained on potato dextrose agar (PDA) at 25°C in the dark. The plate used in the experiment was the result of one passage from a plate grown under identical conditions from a cryogenic stock.

### Experimental design

There are three key points in the fungal life cycle at which termites may detect a terminal infection. The earliest is after conidial germination: as a consequence of the thin, unsclerotised termite cuticle and the limited ability of the individual immune system to encapsulate germ tubes [31], an internal infection can be established within hours of germination (Supplementary Material), and the risks to the colony at this stage may outweigh the benefits of saving an individual on the cusp of infection. The second point is after internal infection has begun and the host has begun to show visible signs of disease [31]. Given the rapidity with which this fungus infects and kills its termite host [32], it is possible that sickness cues such as volatiles or modifications to the cuticular hydrocarbon profile, if present, may take time to produce. The last point at which the colony might detect and respond to a terminal infection is therefore shortly before the termite’s death.

Based on these considerations, we chose to observe the collective responses to a worker termite at one the following four stages of infection: (1) the conidia have attached but not germinated; (2) the conidia have begun to germinate but the host remains healthy; (3) the host is moribund (an internal infection has been established); and (4) the host is near death. To obtain individuals at each stage of infection, termites were treated with a 1 × 10^8^ conidia/mL suspension of *M. anisopliae* or 0.05% Tween 80 as a control, then maintained individually for 2, 12, 15, or 20 hours. These four incubation times were chosen based on the results of a preliminary experiment (Supplementary Material). The reasoning for each is also summarised in Table 1.

**Table 1.**
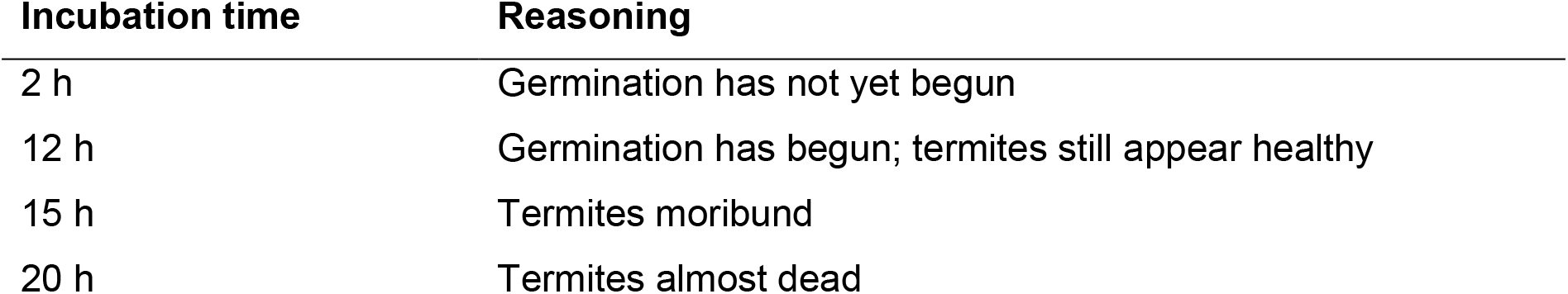
Reasoning for the incubation times chosen in this experiment

| Incubation time | Reasoning |
| --- | --- |
| 2 h | Germination has not yet begun |
| 12 h | Germination has begun; termites still appear healthy |
| 15 h | Termites moribund |
| 20 h | Termites almost dead |

Three colonies were used in the experiment. For each of the four incubation times, there were 24 replicates of the *M. anisopliae* treatment (8 per colony for three colonies) and 12 of the control treatment (4 per colony) to control for the effects of handling and isolation. These were split evenly across two experimental replicates. In the first experimental replicate, the conidia used in the experiment were freshly harvested from half of one PDA plate. In the second experimental replicate, conidia were freshly harvested from the other half.

### Preparation of Petri dish nests

Each Petri dish nest consisted of a Petri dish (94 x 16 mm, without vents), two thick Pall cellulose pads (45.5 mm diameter, 0.9 mm thick), two thin Whatman No. 5 filter paper discs (47 mm diameter, 0.2 mm thick), and one standard glass microscope slide (76 x 26 mm). Each thick cellulose pad was placed on top of a thin filter paper disc. The two paper stacks were then placed side-by-side in the Petri dish, with one of the stacks trimmed on one side to fit. Finally, a glass microscope slide was placed on top (Figure 1). The thickness of the paper stacks (ca. 1.1 mm) was experimentally determined such that when termites dug in the paper under the glass slide, they were able to move freely but had too little space to leave an opaque “ceiling” over their tunnels. The paper was moistened with 3.5 mL tap water.

**Figure 1.**
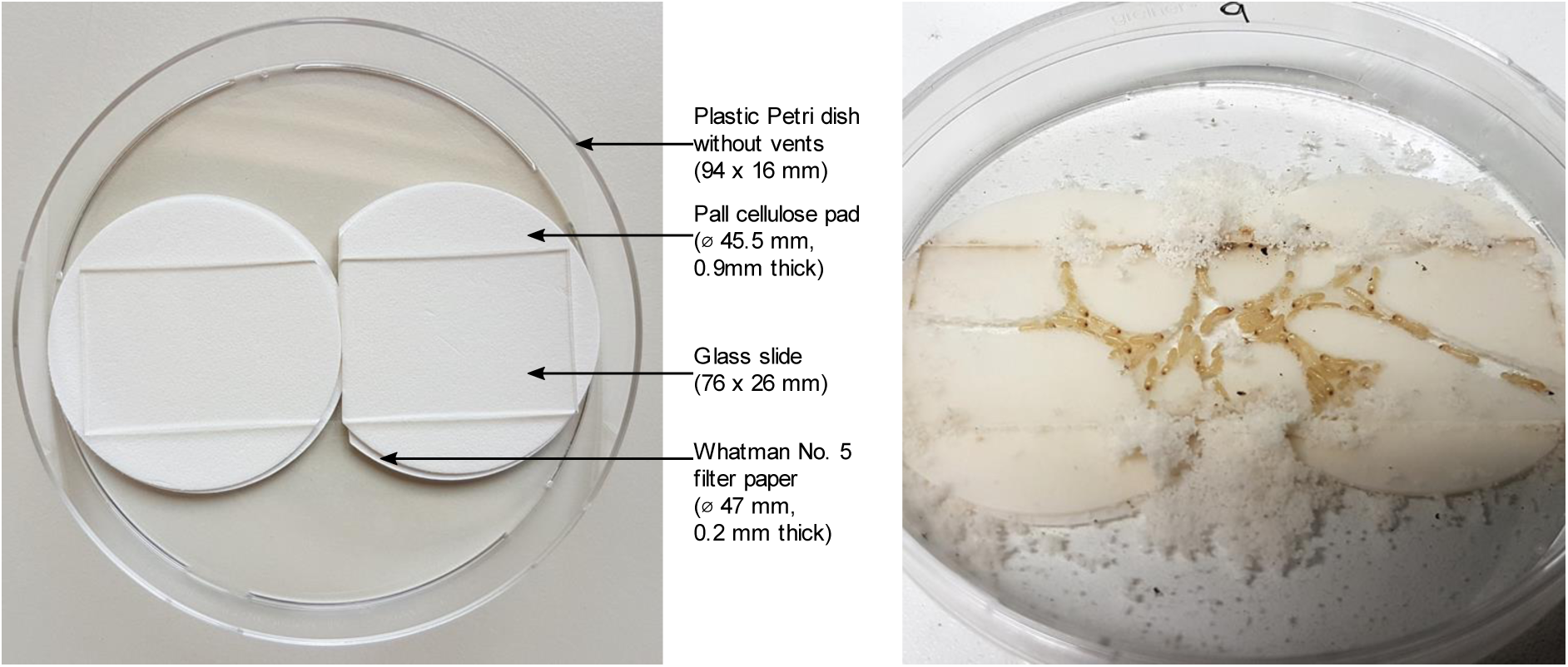
a) Petri dish nest design; b) Petri dish nest after two weeks of termite activity.

Forty-five workers were added to each Petri dish, 1 soldier was added to 20 dishes per colony (10 in the first experimental replicate, 10 in the second), and every dish received 3 representatives of the reproductive caste, with the exception of 7 plates from colony 5, which only received 2 in the second experimental replicate due to difficulty retrieving nymphs from the colony. *R. flavipes* caste ratios vary [33], but workers are by far the dominant caste [19]. The numbers of reproductives and soldiers were taken into account in the statistical analysis, but neither had a significant effect.

In total, each dish contained 48 or 49 termites, not including the focal individual. Becker [30] recommends using a minimum of approximately 50 termites to maintain *R. flavipes* in the lab. In pilot experiments, we confirmed that groups this size could survive for three or more weeks in a Petri dish setup, and that they displayed typical social and hygienic behaviours, including cannibalism and burial. Smaller groups had lower survivorship and sometimes displayed abnormal behaviour, such as leaving corpses uneaten and unburied.

The dishes were sealed with parafilm to prevent desiccation and left in a dark room at 27°C, 70% humidity for two weeks. At least half an hour prior to a behavioural experiment, a cotton swab was used to sweep debris off the glass slide. This was necessary to ensure a clear view into the nest.

### Marking focal termites

Focal termites were marked with Nile blue (AppliChem GmbH, Darmstadt, Germany), a moderately toxic fat-soluble stain that has previously been used to mark termites in behavioural studies [16, 19, 34]. As an internal stain, it cannot be removed and does not interfere with grooming.

Our protocol is a faster version of Evans’ fast marking technique [35]. As termites will swallow any liquid that they are immersed in, we dispensed with his desiccation step and immersed all termites that needed to be stained in 0.025% Nile blue. This is the minimum concentration needed to reliably stain *R. flavipes*.

Large (≥ 4 mm) workers were poured into 2 mL microcentrifuge tubes, one per colony, using a small funnel. Only workers that appeared healthy and active were used. Sufficient 0.025% Nile blue was added to cover them, and they were flicked to mix for 1 minute, then tipped out onto a dry cellulose pad. Initially, all appeared unstained. The termites were transferred to one of three labelled round plastic containers (ca. 52mm inner diameter), one per colony, each lined with a clean cellulose pad moistened with 1 mL tap water and closed with a tight-fitting lid, and then left overnight in a dark room at 27°C, 70% humidity. Only termites that were successfully stained and appeared healthy and active were used in the subsequent experiment. Because the intensity of the colour varied widely, and because of the known toxicity of the stain, termites of different shades were randomly distributed amongst treatments and controls.

### Preparation of conidial suspensions

Conidia were harvested after a minimum of one month of growth. A sterile cotton swab moistened with sterile 0.05% Tween 80 was used to wipe the conidia off the plate and suspend them in sterile 0.05% Tween 80. The suspension was inverted and vortexed to mix, then filtered through a piece of sterile cheese cloth that had been folded to reduce the effective pore size. The filtered conidia were washed by centrifuging for 10 minutes at 5000 g in a centrifuge cooled to 4°C, discarding the supernatant, and resuspending the pellet in sterile 0.05% Tween 80. This step was performed a total of three times.

A BLAUBRAND® Thoma counting chamber (depth 0.1 mm; BRAND, Wertheim, Germany) was used to estimate the concentration of the conidial suspension. Conidial suspensions were adjusted to 1 × 10^8^ conidia/mL with 0.05% Tween 80, aliquoted for ease of use, and used within 48 hours. Suspensions were stored at 4°C when not in use.

To ensure that the conidia were viable, PDA plates were streaked with conidia from an aliquot of the same 1 × 10^8^ conidia/mL suspension used to inoculate the termites. The plates were parafilmed and placed upside-down in the same room as the termites (27°C, 70% humidity). After 21 hours, at least 300 conidia were evaluated for germination at 200 to 400x magnification on one of the plates to calculate the germination rate. A conidium was considered germinated if the length of the germ tube was at least half the diameter of the conidium. For confirmation, at least 100 conidia were evaluated in the same manner on the second plate. The germination rate was ca. 94% in the first experimental replicate and ca. 98% in the second. A germination rate lower than 90% would have indicated a problem with the conidial suspension.

### Inoculation with conidia or 0.05% Tween 80

For the *M. anisopliae* treatment, previously-marked (blue) termites were placed in a round-bottomed 2 mL microcentrifuge tube, then covered with the 1 × 10^8^ conidia/mL suspension to a volume of 42 µL per termite. The tube was flicked to mix for 10 seconds, then poured out onto a dry cellulose pad. Termites that remained inside were tapped out, or, if needed, carefully removed with soft forceps. When the termites had recovered enough to walk, they were transferred one-by-one into separate Petri dishes, each containing a cellulose pad moistened with 1 mL tap water. The dishes were sealed with parafilm to prevent desiccation. Control termites were immersed in sterile 0.05% Tween 80 (42 µL per termite) instead of the conidial suspension and handled in the same way. This inoculation method is a variation on that used by Yanagawa and Shimizu [15].

The *M. anisopliae*-treated and control termites were incubated for 2, 12, 15, or 20 hours at 27°C, 70% humidity before use in the behavioural experiment.

### Behaviour recording

After 2, 12, 15, or 20 hours of incubation, the blue *M. anisopliae*-treated and control focal termites were added individually to the Petri dish nests. All dishes were resealed with parafilm. This took approximately 15 minutes, and the observation period began immediately after the last dish was sealed. Termites that appeared injured or dead at the beginning of an observation period were excluded from the analysis. In total, two replicates of the 2h/*M.a*+ (2 hours of incubation with *M. anisopliae*) treatment, three replicates of the 15h/*M.a*+ (15 hours of incubation with *M. anisopliae*) treatment, and one replicate of the 20h/*M.a*+ (20 hours of incubation with *M. anisopliae*) treatment were excluded due to suspected handling injuries.

Scan sampling [36] was used to observe the interactions between the focal termite and its nestmates within each Petri dish nest. Scans typically took less than 1 minute. They were performed every 5 minutes for a total of 3 hours using a magnifying glass (up to 3x magnification) to better distinguish between similar behaviours and a Samsung S7 smartphone as a digital voice recorder. All observations were made at 27°C, 70% humidity under bright, constant overhead light. As *R. flavipes* are known to respond strongly to vibrational stimuli [37], Petri dishes were not moved or opened after they had been sealed.

States were defined prior to the experiment. We classified behaviours into visually distinguishable, non-overlapping categories with a focus on interactions (and their aftermath) that are relevant to social immunity:

> **Groomed by n**: Focal termite is being groomed by n nestmates with no evidence of biting.
>
> **Bitten**: Focal termite is being bitten by 1 or more nestmates.
>
> **Dismembered**: Focal termite is missing one or more tagmata.
>
> **Dead-ignored**: Focal termite is lying completely motionless, but not buried or dismembered. Nestmates are not interacting with it.
>
> **Not visible**: Focal termite is in a section of the nest where its behaviour and interactions with nestmates cannot be seen.
>
> **Other**: Focal termite is alive, intact, and unburied, but nestmates are not interacting with it.

### Statistical analysis

All statistical analyses were performed using R (version 3.4.3) [38].

#### Grooming

The amount of grooming in each treatment (number of grooming states/total observed states) was compared by fitting a generalised linear mixed model to the data using the glmer function in the package lme4 [39]. Because we were working with proportion data, we used a binomial error structure [40].

The model contained an interaction between incubation time and *M. anisopliae* presence as a fixed effect and two random effects: colony nested within experimental replicate and plate ID. We initially included soldier number and reproductive number as fixed effects, then sequentially removed them during model simplification, using the anova function to ascertain if the removal of a parameter would lead to a significant change in deviance and to perform likelihood ratio test comparisons. The final model was tested for overdispersion using the dispersion_glmer function in the package blmeco [41]. A scale parameter between 0.75 and 1.4 indicates no overdispersion: for this model, it was 1.004. All post hoc pairwise comparisons were performed using the glht function from the multcomp package [42] with Tukey correction.

To analyse grooming intensity, we used glmer to fit a generalised linear mixed model to the data with the total number of groomers in each replicate as the response variable and a Poisson error structure. Two replicates (one in the 2 hour control treatment, one in the 15 hour control treatment) were excluded from the analysis because no grooming states were observed. The model contained an interaction between incubation time and *M. anisopliae* presence as a fixed effect and two random effects: colony nested within experimental replicate and plate ID. The log of the number of grooming states was used as an offset. We initially included soldier number and reproductive number as fixed effects, then sequentially removed them during model simplification, using anova as above to compare models. The scale parameter of the final model was 0.861. All post hoc pairwise comparisons were performed using glht with Tukey correction.

#### Cannibalism

“Bitten” and “dismembered” states were combined into a single “cannibalism” state. We modelled the onset of cannibalistic behaviour using survival curves. The data were plotted using survfit from the survival package [43, 44] and ggsurvplot from the survminer package [45]. We used a mixed effects Cox model (coxme from the coxme package [46]) to compare the curves.

The model contained an interaction between incubation time and *M. anisopliae* presence as a fixed effect, and colony nested within experimental replicate as a random effect. We initially included soldier number and reproductive number as fixed effects, then sequentially removed them during model simplification, using the anova function as above to compare models. In the survival curve analysis, all control data was initially right-censored; in order to fit a mixed effects Cox model to the data, it was necessary to uncensor one arbitrarily-selected control replicate from each incubation time treatment following Tragust et al. [47]. The glht function was used to perform post-hoc pairwise comparisons with Tukey correction.

## Results

### Patterns of behaviour

Focal termites were visible throughout the observation period, and there were no instances of focal termite corpses being ignored. Behavioural patterns (Figure 2) in the control treatments (2h/*M.a*-, 12h/*M.a*-, 15h/*M.a*-, 20h/*M.a*-) were broadly similar, dominated by states in the “other” category, with low levels of grooming and no cannibalism or burial states. The majority of states in 2h/*M.a*+ were in the “other” category, but grooming was elevated over the control. 12h/*M.a*+ was dominated by high levels of grooming that slowly decreased over the observation period, while “other” states slowly increased. Cannibalism was observed, but primarily in the last half hour of the observation period. No burial states were recorded. Both 15h/*M.a*+ and 20h/*M.a*+ were characterised by high levels of intense grooming immediately after the focal termites were introduced. Cannibalism began shortly thereafter, increasing more rapidly in 20h/*M.a*+ and completely replacing grooming before the end of the observation period. Burial was observed in both treatments toward the end of the observation period, but only in a small proportion of states.

**Figure 2.**
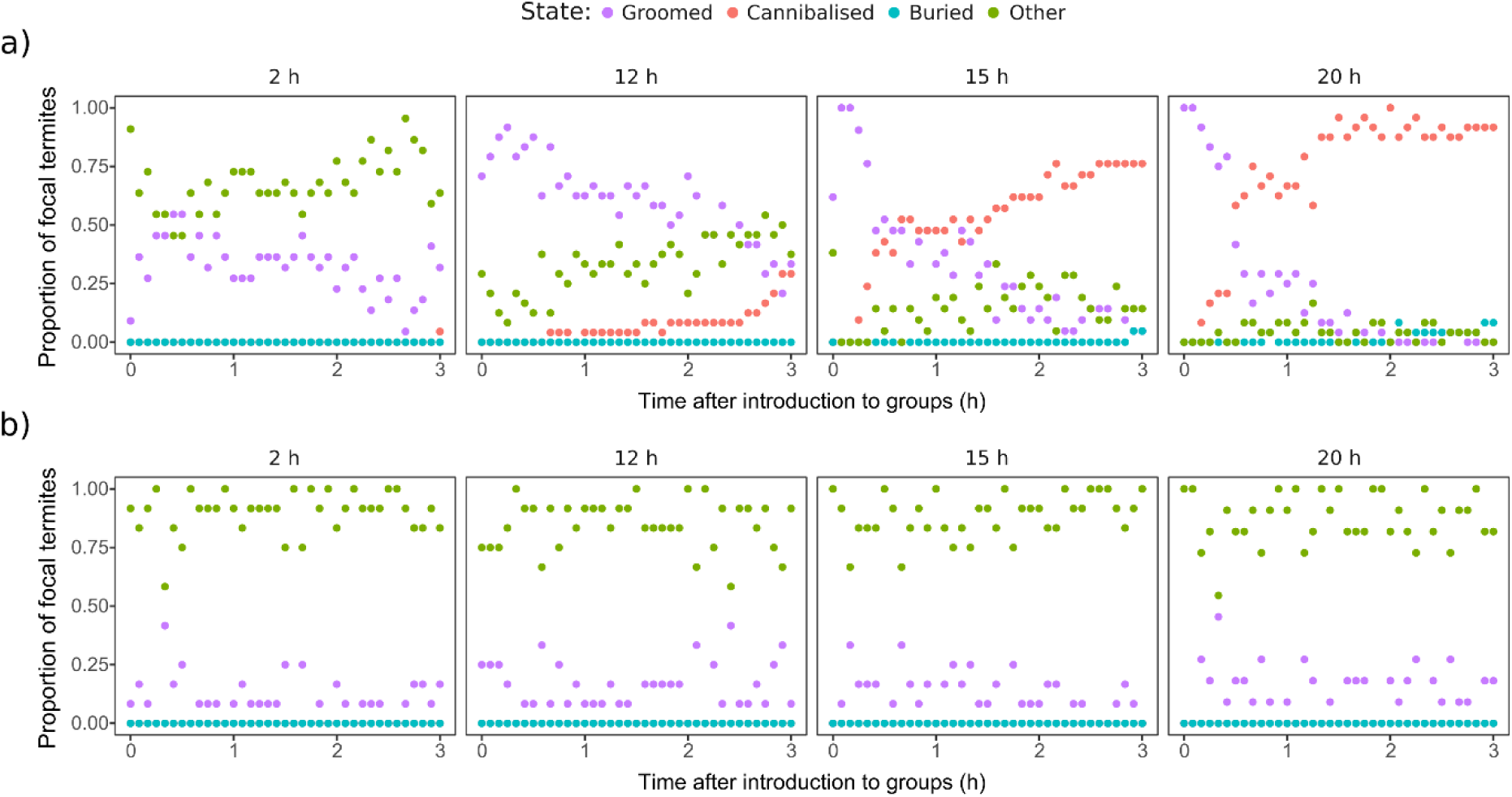
Patterns of behaviour over time during the three-hour observation period for a) *M. anisopliae* and b) control treatments. Each point represents the proportion of focal termites that were observed in a given state during that scan. When more than one state was present at the same proportion (0.50 or 0.00), the points overlap.

### Grooming

The proportion of states classified as grooming in the 12h/*M.a*+ treatment was significantly elevated over all other *M. anisopliae* treatments (12h/*M.a*+ vs. 2h/*M.a*+ z=5.533 p<0.001; 15h/*M.a*+ vs. 12h/*M.a*+ z=-5.511 p<0.001; 20h/*M.a*+ vs. 12h/*M.a*+ z=-7.717 p<0.001) (Figure 3, Table S1). Grooming was significantly elevated over the controls in all *M. anisopliae* treatments except 20h/*M.a*+, which was only significantly different from 2h/*M.a*-(*M. anisopliae* treatments vs. corresponding controls: 2h/*M.a*+ vs. 2h/*M.a*-z=4.844 p<0.001; 12h/*M.a*+ vs. 12h/*M.a*-z=7.834 p<0.001; 15h/*M.a*+ vs. 15h/*M.a*-z=4.417 p<0.001). The controls (2h/*M.a*-, 12h/*M.a*-, 15h/*M.a*-, 20h/*M.a*-) were not significantly different from each other. No significant difference was observed between 2h/*M.a*+, 15h/*M.a*+, and 20h/*M.a*+. Low levels of grooming in 15h/*M.a*+ and 20h/*M.a*+ correspond with a high proportion of cannibalism states in both treatments (Figure 2, Figure 5).

**Figure 3.**
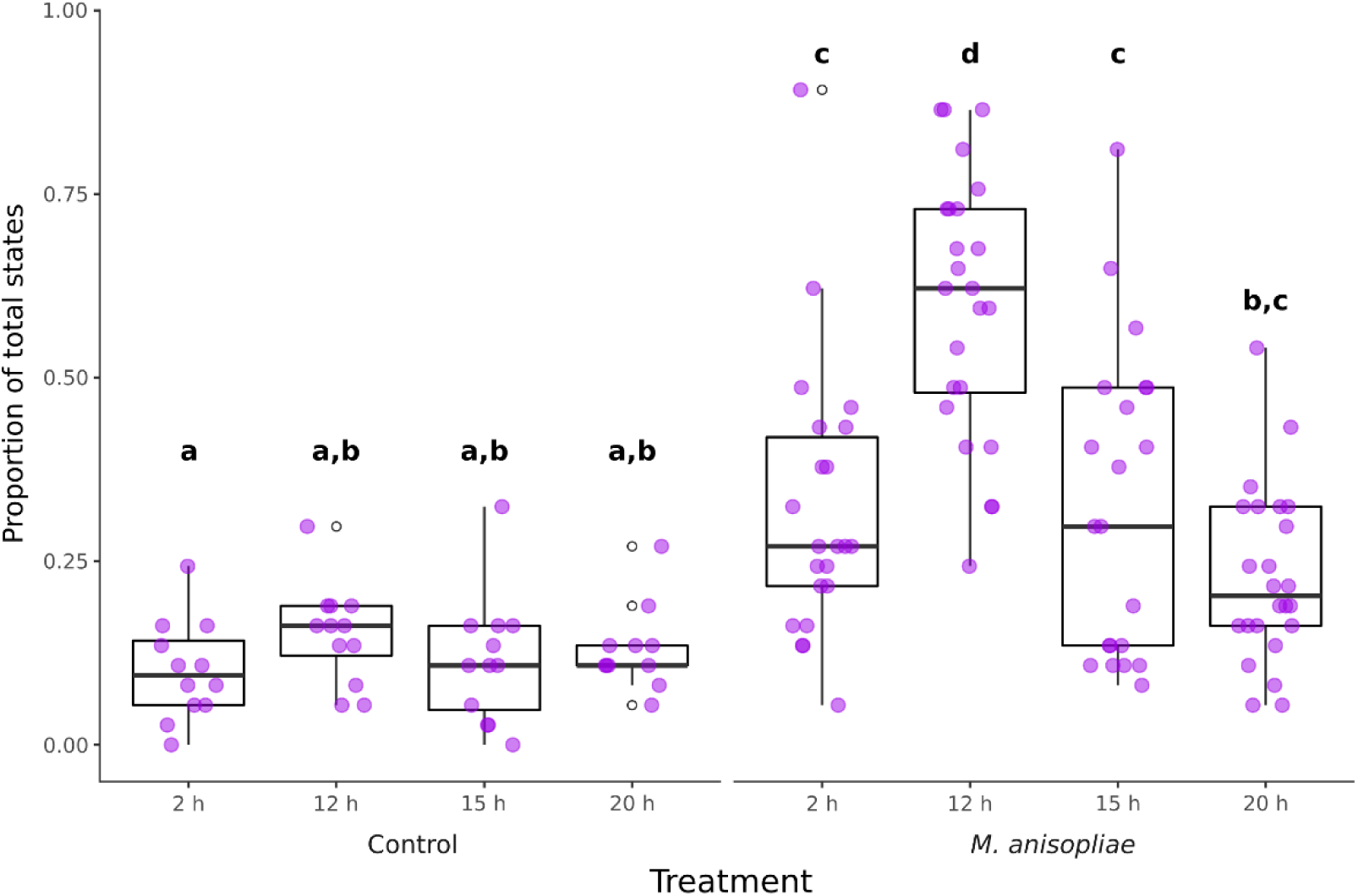
Grooming as a proportion of total states. Treatments marked with different letters were significantly different (Table S1). Lower and upper hinges correspond to first and third quartiles, the upper whisker extends to the largest value if it is no greater than 1.5 times the inter-quartile rage from the hinge, and the lower whisker extends to the smallest value if it is no smaller than 1.5 times the inter-quartile range from the hinge.

Only workers were observed grooming the focal termites, and grooming was visibly more intense, involving a significantly higher number of groomers, in 12h/*M.a*+, 15h/*M.a*+, and 20h/*M.a*+ (12h/*M.a*+ vs. 2h/*M.a*+ z=3.299 p=0.01834; 15h/*M.a*+ vs. 2h/*M.a*+ z=7.915 p<0.001; 15h/*M.a*+ vs. 12h/*M.a*+ z=5.757 p<0.001; 20h/*M.a*+ vs. 2h/*M.a*+ z=8.077 p<0.001; 20h/*M.a*+ vs. 12h/*M.a*+ z=5.873 p<0.001) (Figure 4, Table S2). 2h/*M.a*+ was not significantly different from any of the controls, and 15h/*M.a*+ and 20h/*M.a*+ were not significantly different from each other. Non-focal termites in treatments with more intense grooming were frequently observed to engage in vibratory displays (jittering), a known pathogen alarm response [11, 48]; however, our sampling method, which focused on direct interactions with the focal individual, precluded analysis of this behaviour.

**Figure 4.**
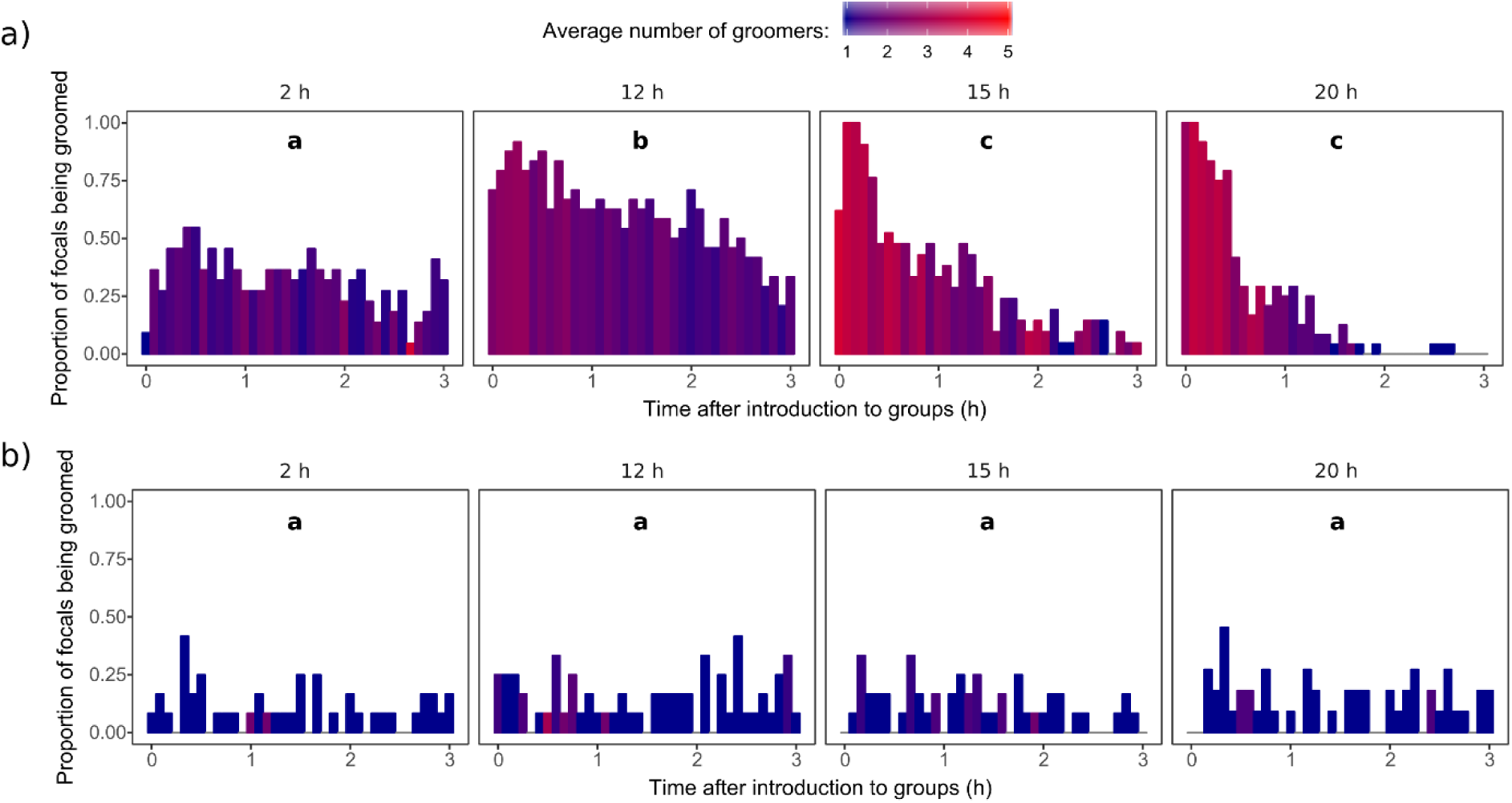
Proportion of focal termites in the (a) *M. anisopliae* or (b) control treatments observed being groomed by nestmates in each scan (as in Figure 2), with the fill colour representing the average number of groomers involved. Different letters correspond to significant differences in the overall number of groomers after the number of grooming states is taken into account (Table S2).

### Cannibalism

The probability of remaining unharmed during the observation period was significantly different from the controls, which experienced no cannibalism, in the 15h/*M.a*+ (z=3.958 p=0.00167) and 20h/*M.a*+ (z=4.809 p<0.001) treatments, but not in 2h/*M.a*+ or 12h/*M.a*+ (Table S3). The 15h/*M.a*+ and 20h/*M.a*+ treatments differed significantly from each other (z=3.006, p=0.04437), as well as from 2h/*M.a*+ and 12h/*M.a*+ (15h/*M.a*+ vs. 2h/*M.a*+ z=4.550 p<0.001; 15h/*M.a*+ vs. 12h/*M.a*+ z=5.233 p<0.001; 20h/*M.a*+ vs. 2h/*M.a*+ z=5.469 p<0.001; 20h/*M.a*+ vs. 12h/*M.a*+ z=6.745 p<0.001). This difference was characterised by an earlier onset of cannibalism in 20h/*M.a*+ than in 15h/*M.a*+ (Figure 5). In all but two cases, the first cannibalism-related state recorded was biting. In those two exceptions, both in 15h/*M.a*+, it was dismemberment. The previous scan recorded intense grooming of the focal individual by 5 to 6 groomers: it is possible that that state was misidentified, or that biting began between scans. Cannibalism was performed primarily by workers, but on two occasions, a brachypterous neotenic was observed to also partake.

**Figure 5.**
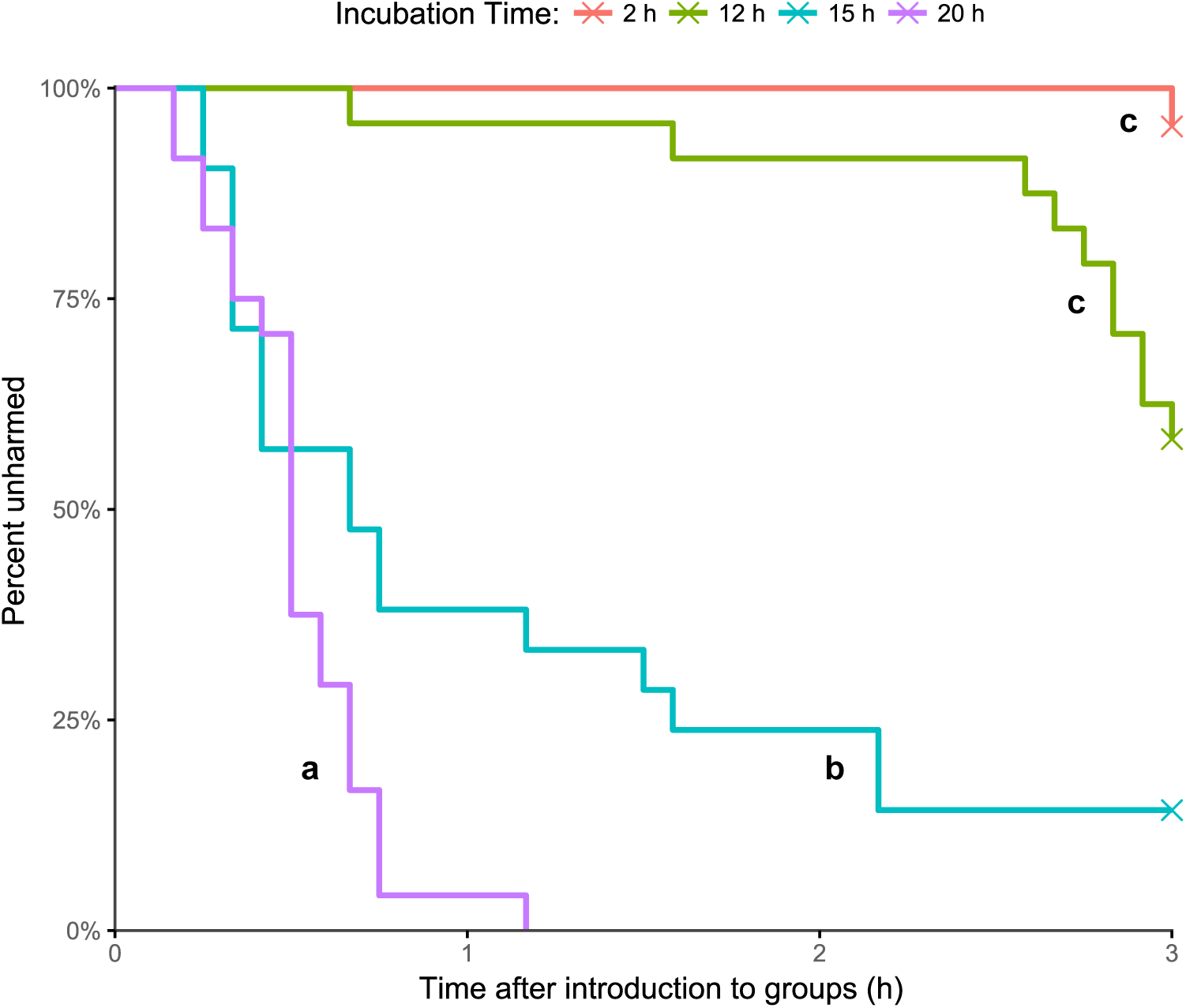
Percentage of *M. anisopliae-treated* focal termites that remained unharmed (not bitten or dismembered) over the three-hour observation period. X’s indicate the presence of right-censored data (i.e. focal termites that were not harmed during the observation period). Treatments marked with the same letter were not significantly different, and neither “c” treatment (2h/*M.a*+, 12h/*M.a*+) was significantly different from the controls (not shown; all control individuals remained unharmed throughout the observation period) (Table S3).

### Burial

Burial was observed in four plates: one 15h/*M.a*+ replicate and three 20h/*M.a*+ replicates. This is too few for meaningful statistical analysis. In each case, the focal termite appeared to be alive but moribund and largely immobile at the beginning of the burial process. This immobility was caused by nestmates in one 20h/*M.a*+ replicate: the legs were first bitten off, and the maimed termite was left for approximately half an hour before burial began.

There was no sudden switch from grooming or cannibalism to burial. In one 20h/*M.a*+ replicate, the focal termite was initially groomed, then bitten, then had a piece of paper placed on top of it (burial), then groomed again for half an hour, during which time the paper was removed, then bitten again. Burial did not resume, and the termite was eventually dismembered.

## Discussion

Our results demonstrate that *R. flavipes* colonies employ different collective immune defence strategies at different stages of infection with *M. anisopliae*. Before conidia germinate, the social immune response is dominated by grooming; however, contrary to our first hypothesis, levels of grooming rise significantly after conidial germination, and it becomes visibly more intense. Contrary to our second hypothesis, the onset of cannibalistic behaviour coincides with the stage of infection in which the termite becomes moribund, with a more rapid switch to cannibalism at later stages. All cannibalised individuals were eaten alive. This is consistent with observations by Rosengaus and Traniello [3], who observed that termites were usually cannibalised when near death, but contradicts Strack [24], who observed more “agonism” toward healthy individuals that had been thickly dusted with conidia. Burial was rarely observed, reinforcing the view that termites preferentially eliminate sick individuals through cannibalism [3, 23].

The unexpectedly low levels of grooming observed before conidial germination may be explained by their weak attachment to the cuticle: since most conidia can be removed within hours by relatively few individuals [26], there may be no reason to divert resources away from other colony functions or endanger additional members of the colony. The effectiveness of allogrooming, even at the observed low intensity, can also be seen in survivorship studies [12]. Increased levels of grooming after germination, then, could be linked to increased physical difficulty removing fungal material, especially after germ tube penetration.

This explanation is unsatisfying, because the longer a pathogen persists on or in members of a colony, the more we would expect it to affect colony fitness. Conidia-exposed, non-moribund individuals are mobile and can transfer conidia to many colony members [13, 14], all of which would need to be groomed by workers that could otherwise be performing other tasks. Should the infection progress to the next stage, the risk to the colony would increase significantly. This should favour early “clearance” of the infection from the colony via aggregation and intense grooming of conidia-exposed individuals, but that is not what we observed.

A second possibility is that the fungus-associated molecules that stimulate grooming (e.g. the fungal “odour” [49]) are partly masked, or present in lower quantities, before germination. Based on response threshold models of division of labour in social insects [50], even partial masking would result in a weaker collective grooming response with fewer participating workers. This need not be a specific adaptation to evade termite social immunity, nor would we expect it in a generalist entomopathogen. Masking of immunogenic components of the fungal cell wall before (but not after) germination has previously been reported in an opportunistic human pathogen, *Aspergillus fumigatus* Fresenius [51]. Should this prove to be the case in *M. anisopliae*, it could be harnessed to develop strains with higher epizootic potential.

In contrast to grooming, in which fungal factors appear to be the primary trigger, the strong temporal correlation between moribundity and cannibalistic behaviour suggests that the host plays a central role in its own sacrifice. Focal termites appeared healthy at 12 hours and moribund (a reliable sign of internal infection [31]) at 15, and cannibalism was only prevalent in the 15h/*M.a*+ and 20h/*M.a*+ treatments. Even in the 12h/*M.a*+ treatment, which was not significantly different from the control, there was an uptick in cannibalism in the last half hour of the observation period, i.e. at approximately 14.5 hours post-exposure. With the caveat that this is a correlation, and that moribundity could coincide with some fungus-derived stimulus reaching the necessary threshold for cannibalism, the hypothesis that sick individuals might flag themselves for destruction is supported by research in the social Hymenoptera. Ant pupae “advertise” the presence of an internal infection through modified cuticular hydrocarbon profile [4], and aggressive behaviour was observed toward adults at the same stage of infection [10]; however, more work will be required to determine whether the social Blattodea and the social Hymenoptera have independently evolved separate mechanisms to identify fatally ill colony members, or if they have separately co-opted evolutionarily conserved sickness cues for social immune defence.

## Conclusion

We have demonstrated that termites can deploy different collective immune defences when confronted with a worker at different stages of infection with an entomopathogenic fungus. Whereas grooming is favoured earlier in the infectious process, moribund individuals are readily sacrificed to protect the colony. Cannibalism appears to be triggered by some factor associated with moribundity: what this might be remains unclear. Paradoxically, the termites did not display a robust social immune response at the earliest stages, when conidia had not yet germinated, although grooming was somewhat elevated. This may indicate that the ungerminated fungus is less visible to the “social immune system” of the colony, but this hypothesis remains to be tested.

This study adds to the body of knowledge surrounding termite social immunity and sheds light on how colonies resist fungal disease and regulate destructive immune behaviours. By dividing the infection into stages [29] and studying how the social immune response differs over time, we can better understand how termites, and insects in general, defend their colonies from disease.

## Acknowledgements

We thank R. Plarre and J. Rolff for discussion and support, as well as A. Herrmann, S. He, and especially Y. de Laval for assistance in the laboratory.

## References

1. Cremer S., Armitage S.A.O., Schmid-Hempel P. 2007 Social Immunity. Curr Biol 17(16), 693–702. (doi:10.1016/j.cub.2007.06.008).

2. Page P., Lin Z., Buawangpong N., Zheng H., Hu F., Neumann P., Chantawannakul P., Dietemann V. 2016 Social apoptosis in honey bee superorganisms. Sci Rep 6, 27210. (doi:10.1038/srep27210).

3. Rosengaus R.B., Traniello J.F.A. 2001 Disease susceptibility and the adaptive nature of colony demography in the dampwood termite Zootermopsis angusticollis. Behav Ecol Sociobiol 50(6), 546–556. (doi:10.1007/s002650100394).

4. Pull C.D., Ugelvig L.V., Wiesenhofer F., Grasse A.V., Tragust S., Schmitt T., Brown M.J.F., Cremer S. 2018 Destructive disinfection of infected brood prevents systemic disease spread in ant colonies. eLife 7, e32073. (doi:10.7554/eLife.32073.001).

5. Richard F.-J.J., Aubert A., Grozinger C.M. 2008 Modulation of social interactions by immune stimulation in honey bee, *Apis mellifera*, workers. BMC Biol 6, 50. (doi:10.1186/1741-7007-6-50).

6. Baracchi D., Fadda A., Turillazzi S. 2012 Evidence for antiseptic behaviour towards sick adult bees in honey bee colonies. J Insect Physiol 58(12), 1589–1596. (doi:10.1016/j.jinsphys.2012.09.014).

7. Nazzi F., Della Vedova G., D’Agaro M. 2004 A semiochemical from brood cells infested by *Varroa destructor* triggers hygienic behaviour in *Apis mellifera*. Apidologie 35(1), 65–70. (doi:10.1051/apido:2003065).

8. Spivak M., Downey D.L. 1998 Field Assays for Hygienic Behavior in Honey Bees (Hymenoptera: Apidae). J Econ Entomol 91(1), 64–70. (doi:10.1093/jee/91.1.64).

9. Rosenkranz P., Tewarson N.C., Singh A., Engels W. 2015 Differential hygienic behaviour towards *Varroa jacobsoni* in capped worker brood of *Apis cerana* depends on alien scent adhering to the mites. J Apic Res 32(2), 89–93. (doi:10.1080/00218839.1993.11101292).

10. Leclerc J.B., Detrain C. 2016 Ants detect but do not discriminate diseased workers within their nest. Sci Nat 103(7-8), 70. (doi:10.1007/s00114-016-1394-8).

11. Myles T.G. 2002 Alarm, aggregation, and defense by Reticulitermes flavipes in response to a naturally occurring isolate of Metarhizium anisopliae. Sociobiology 40(2), 243–255.

12. Rosengaus R.B., Maxmen A.B., Coates L.E., Traniello J.F.A. 1998 Disease resistance: a benefit of sociality in the dampwood termite *Zootermopsis angusticollis* (Isoptera: Termopsidae). Behav Ecol Sociobiol 44(2), 125–134. (doi:10.1007/s002650050523).

13. Liu L., Li G., Sun P., Lei C., Huang Q. 2015 Experimental verification and molecular basis of active immunization against fungal pathogens in termites. Sci Rep 5, 15106. (doi:10.1038/srep15106).

14. Kramm K.R., West D.F., Rockenbach P.G. 1982 Termite pathogens: Transfer of the entomopathogen *Metarhizium anisopliae* between *Reticulitermes* sp. termites. J Invertebr Pathol 40(1), 1–6. (doi:10.1016/0022-2011(82)90029-5).

15. Yanagawa A., Shimizu S. 2007 Resistance of the termite, *Coptotermes formosanus* Shiraki to *Metarhizium anisopliae* due to grooming. BioControl 52(1), 75–85. (doi:10.1007/s10526-006-9020-x).

16. Yanagawa A., Fujiwara-Tsujii N., Akino T., Yoshimura T., Yanagawa T., Shimizu S. 2011 Behavioral changes in the termite, *Coptotermes formosanus* (Isoptera), inoculated with six fungal isolates. J Invertebr Pathol 107(2), 100–106. (doi:10.1016/j.jip.2011.03.003).

17. Myles T.G. 2002 Laboratory studies on the transmission of *Metarhizium anisopliae* in the eastern subterranean termite, *Reticulitermes flavipes* (Isoptera: Rhinotermitidae), with a method for applying appropriate doses of conidia to trapped termites for release. Sociobiology 40(2), 265–276.

18. Logan J.W.M., Cowie R.H., Wood T.G. 1990 Termite (Isoptera) control in agriculture and forestry by non-chemical methods: a review. Bull Entomol Res 80(3), 309–330. (doi:10.1017/s0007485300050513).

19. Chouvenc T., Su N.-Y., Elliott M.L. 2008 Interaction between the subterranean termite *Reticulitermes flavipes* (Isoptera: Rhinotermitidae) and the entomopathogenic fungus *Metarhizium anisopliae* in foraging arenas. J Econ Entomol 101(3), 885–893. (doi:10.1093/jee/101.3.885).

20. Rosengaus R.B., Traniello J.F.A., Bulmer M.S. 2010 Ecology, Behavior and Evolution of Disease Resistance in Termites. In *Biology of Termites: a Modern Synthesis* (eds. Bignell D.E., Roisin Y., Lo N.), pp. 165–191. Dordrecht, Springer.

21. Sun Q., Haynes K.F., Zhou X., Ayasse M. 2017 Dynamic changes in death cues modulate risks and rewards of corpse management in a social insect. Funct Ecol 31(3), 697–706. (doi:10.1111/1365-2435.12754).

22. Neoh K.-B., Yeap B.-K., Tsunoda K., Yoshimura T., Lee C.-Y. 2012 Do termites avoid carcasses? Behavioral responses depend on the nature of the carcasses. PLoS One 7(4), e36375. (doi:10.1371/journal.pone.0036375).

23. Chouvenc T., Su N.-Y. 2012 When subterranean termites challenge the rules of fungal epizootics. PLoS One 7(3), e34484. (doi:10.1371/journal.pone.0034484).

24. Strack B.H. 1998 The role of social behaviour of *Reticulitermes flavipes* (Kollar) (Isoptera: Rhinotermitidae) in defence against the fungal pathogen *Metarhizium ansiopliae* (Metschikoff) Sorokin (Deuteromycotina: Hyphomycetes), National Library of Canada = Bibliothèque nationale du Canada.

25. Yanagawa A., Yokohari F., Shimizu S. 2009 The role of antennae in removing entomopathogenic fungi from cuticle of the termite, *Coptotermes formosanus*. J Insect Sci 9, 6. (doi:10.1673/031.009.0601).

26. Yanagawa A., Yokohari F., Shimizu S. 2010 Influence of fungal odor on grooming behavior of the termite, *Coptotermes formosanus*. J Insect Sci 10, 141. (doi:10.1673/031.010.14101).

27. Chouvenc T., Su N.-Y., Robert A. 2009 Inhibition of *Metarhizium anisopliae* in the alimentary tract of the eastern subterranean termite *Reticulitermes flavipes*. J Invertebr Pathol 101(2), 130–136. (doi:10.1016/j.jip.2009.04.005).

28. Hänel H. 1982 The life cycle of the insect pathogenic fungus *Metarhizium anisopliae* in the termite *Nasutitermes exitiosus*. Mycopathologia 80(3), 137–145. (doi:10.1007/bf00437576).

29. Hall M.D., Bento G., Ebert D. 2017 The Evolutionary Consequences of Stepwise Infection Processes. Trends Ecol Evol 32(8), 612–623. (doi:10.1016/j.tree.2017.05.009).

30. Becker G. 1969 Rearing of Termites and Testing Methods Used in the Laboratory. I. Biology of Termites (eds. Krishna K., Weesner F.M.), pp. 351–385. New York, Academic Press.

31. Chouvenc T., Su N.-Y., Robert A. 2009 Cellular encapsulation in the eastern subterranean termite, *Reticulitermes flavipes* (Isoptera), against infection by the entomopathogenic fungus *Metarhizium anisopliae*. J Invertebr Pathol 101(3), 234–241. (doi:10.1016/j.jip.2009.05.008).

32. Kramm K.R., West D.F. 1982 Termite pathogens: Effects of ingested *Metarhizium, Beauveria*, and *Gliocladium* conidia on worker termites (*Reticulitermes* sp.). J Invertebr Pathol 40(1), 7–11. (doi:10.1016/0022-2011(82)90030-1).

33. Gao Q., Bidochka M.J., Thompson G.J. 2012 Effect of group size and caste ratio on individual survivorship and social immunity in a subterranean termite. Acta Ethol 15(1), 55–63. (doi:10.1007/s10211-011-0108-7).

34. Traniello J.F., Rosengaus R.B., Savoie K. 2002 The development of immunity in a social insect: evidence for the group facilitation of disease resistance. Proc Natl Acad Sci U S A 99(10), 6838–6842. (doi:10.1073/pnas.102176599).

35. Evans T.A. 2000 Fast marking of termites (Isoptera: Rhinotermitidae). Sociobiology 35(3), 517–523.

36. Altmann J. 1974 Observational Study of Behavior: Sampling Methods. Behaviour 49(3), 227–267. (doi:10.1163/156853974X00534).

37. Hertel H., Hanspach A., Plarre R. 2011 Differences in Alarm Responses in Drywood and Subterranean Termites (Isoptera: Kalotermitidae and Rhinotermitidae) to Physical Stimuli. J Insect Behav 24(2), 106–115. (doi:10.1007/s10905-010-9240-x).

38. R Core Team 2017 R: A Language and Environment for Statistical Computing. Vienna, Austria: R Foundation for Statistical Computing. https://www.R-project.org/.

39. Bates D., Mächler M., Bolker B., Walker S. 2015 Fitting Linear Mixed-Effects Models Using lme4. J Stat Softw 67. (doi:10.18637/jss.v067.i01).

40. Crawley M.J. 2015 Statistics: An introduction using R (2nd Edition). Chichester, John Wiley & Sons, Ltd.

41. Korner-Nievergelt F., Roth T., von Felten S., Guélat J., Almasi B., Korner-Nievergelt P. 2015. Bayesian data analysis in ecology using linear models with R, BUGS and Stan. London, Elsevier.

42. Hothorn T., Bretz F., Westfall P. 2008 Simultaneous inference in general parametric models. Biom J 50(3), 346–363. (doi:10.1002/bimj.200810425).

43. Therneau T.M. 2015 A Package for Survival Analysis in S. Version 2.38. https://CRAN.R-project.org/package=survival.

44. Therneau T.M., Grambsch P.M. 2000 Modeling Survival Data: Extending the Cox Model. New York, Springer.

45. Kassambara A., Kosinski M. 2017 survminer: Drawing Survival Curves using ‘ggplot2’. R package version 0.4.1. https://CRAN.R-project.org/package=survminer.

46. Therneau T.M. 2018 coxme: Mixed Effects Cox Models. R package version 2.2-7. https://CRAN.R-project.org/package=coxme.

47. Tragust S., Ugelvig L.V., Chapuisat M., Heinze J., Cremer S. 2013 Pupal cocoons affect sanitary brood care and limit fungal infections in ant colonies. BMC Evol Biol 13, 225. (doi:10.1186/1471-2148-13-225).

48. Rosengaus R.B., Jordan C., Lefebvre M.L., Traniello J.F.A. 1999 Pathogen Alarm Behavior in a Termite: A New Form of Communication in Social Insects. Naturwissenschaften 86(11), 544–548. (doi: 10.1007/s001140050672)

49. Yanagawa A., Fujiwara-Tsujii N., Akino T., Yoshimura T., Yanagawa T., Shimizu S. 2011 Musty odor of entomopathogens enhances disease-prevention behaviors in the termite *Coptotermes formosanus*. J Invertebr Pathol 108(1), 1–6. (doi:10.1016/j.jip.2011.06.001).

50. Beshers S.N., Fewell J.H. 2001 Models of division of labor in social insects. Annu Rev Entomol 46(1), 413–440. (doi:10.1146/annurev.ento.46.1.413).

51. Aimanianda V., Bayry J., Bozza S., Kniemeyer O., Perruccio K., Elluru S.R., Clavaud C., Paris S., Brakhage A.A., Kaveri S.V., et al. 2009 Surface hydrophobin prevents immune recognition of airborne fungal spores. Nature 460, 1117–1121. (doi:10.1038/nature08264).

