## Supplementary Materials for "When to care and when to kill: termites shape their collective response based on stage of infection"

### Supplementary Material

#### Germination Experiment

##### Methods

For each of 8 incubation times (0, 10, 12, 16, 20, 24, 48, and 72 h post-exposure), 5 *Reticulitermes flavipes* workers from the same colony (Colony F; collected in Soulac-sur-Mer, France in 2015; maintained in opaque plastic boxes containing damp soil and wood at 26°C, 84% humidity in the dark) were chilled in a plastic Petri dish on ice, then individually immersed in 15µL drops of a 1 x $10^8$ conidia/mL *Metarhizium anisopliae* suspension pipetted directly on top of them. The termites were placed on a dry cellulose pad to absorb excess liquid, then transferred to individual wells of a six well plate, each containing one quarter of a Pall cellulose pad moistened with 300 µL water. The plates were incubated at 27°C under low light conditions in a box with wet paper towels to keep humidity high. An equivalent number of controls were briefly immersed in 0.05% Tween 80 and incubated individually under the same conditions.

At each time point, *M. anisopliae*-exposed termites were preserved in 70% ethanol. After all had been preserved in this manner, the termites were dehydrated in increasing ethanol concentrations (30 min 90% ethanol, 2x 30 min 100% ethanol, 2x 30 min anhydrous 100% ethanol), critical point dried using $CO_2$ (12 medium changes; CPD 030, BALTEC, Balzers, Liechtenstein), sputtered with gold in a SCD 040 (Balzers, Liechtenstein), and then imaged with a scanning electron microscope (SEM; FEI Quanta 200, FEI Deutschland GmbH, Frankfurt/Main, Germany). The germination rate was calculated by evaluating at least 100 conidia on the termite head according to the following criteria: a conidium was considered germinated if the germ tube was longer than half the diameter of the conidium. Two termites, one in the 12 h incubation group and one in the 20 h incubation group, were excluded from further analysis because they had too few conidia adhering to their cuticles to calculate a germination rate. After 48 h, it was no longer possible to evaluate germination due to dense fungal and bacterial growth on the termite cuticle (Figure S2).

To evaluate germination in vitro, 5 PDA plates were streaked with the same conidial suspension, parafilmed, and incubated at 27°C under low light. At 0, 10, 12, 16, 20, and 24 h after inoculation, at least 100 conidia per plate were evaluated for germination at 200-400x using a phase contrast microscope. After 24 h, dense hyphal growth made it impossible to calculate a germinate rate.

Means and 95% confidence intervals were calculated and plotted using R (version 3.4.3) [1].

#### Results

Germination was first observed at 10 hours (Figure S1, Figure S3); the earliest time reported in the literature is 12 hours after exposure [2, 3]. In vitro, under ideal conditions, germination can begin as early as 5 hours after inoculation [4], but it is known to be delayed on arthropod cuticles [5].

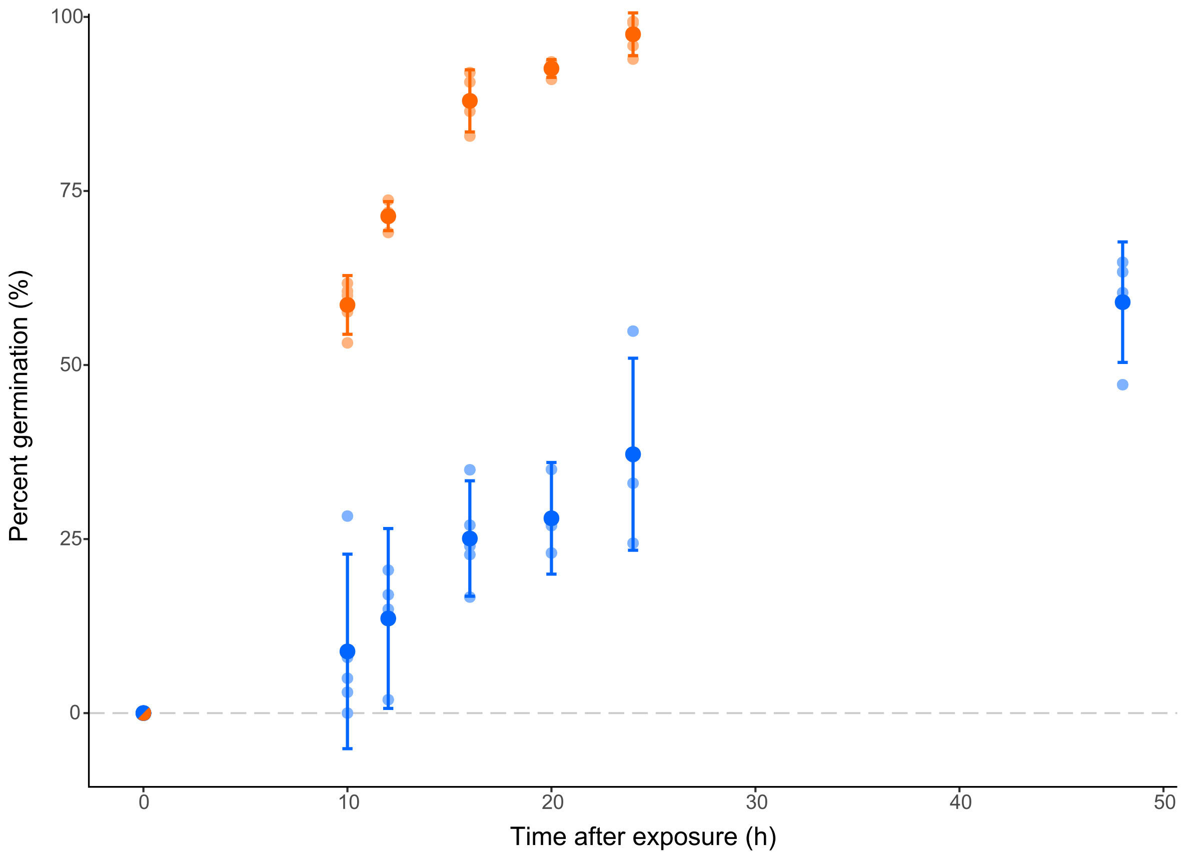


**Figure S1. Germination on PDA plates (orange; the steeper curve) and *Reticulitermes flavipes* worker cuticles (blue). Larger dots represent means, smaller dots are individual data points, and bars show 95% confidence intervals.**

We observed *M. anisopliae*-treated termites showing signs of illness (moribund; when grabbed with forceps, they would bite rather than try to run) as early as 15 hours after exposure to the conidial suspension. The timing of this behavioural change, which correlates with the presence of an internal infection [3] and is consistent with an older high dose study in which moribundity was observed at 16 hours [6], indicates that penetration must begin shortly after germination. Our results contrast with prior fungal life cycle studies which report that penetration occurs 24-36 hours after exposure [3]. The discrepancy may be the result of difficulty spotting penetration: when reviewing SEM images, we were unable to reliably distinguish between germ tubes that had or had not penetrated. Appressoria were occasionally observed at 12 hours after exposure (Figure S4), but as the infection peg, if present, would be underneath the appressorium and hidden from view, we could not say whether penetration had already begun.

At 20 hours, the termites were still alive, but very lethargic. Death occurred between 24 and 48 hours. Germination on the cuticle, which was consistently lower than germination on PDA, continued even after the termite's death from the infection, as did hyphal growth (Figure S5). All controls, with the exception of one that died from an apparent handling injury, were alive and healthy 1 week after the beginning of the experiment.

### References


1. R Core Team 2017 R: A Language and Environment for Statistical Computing. Vienna, Austria: R Foundation for Statistical Computing. https://www.R-project.org/.
2. Moino Jr A., Alves S.B., Lopes R.B., Neves P.M.O.J., Pereira R.M., Vieira S.A. 2002 External development of the entomopathogenic fungi *Beauveria bassiana* and *Metarhizium anisopliae* in the subterranean termite *Heterotermes tenuis*. *Scientia Agricola* **59**(2), 267-273. (doi:10.1590/s0103-90162002000200010).
3. Chouvenc T., Su N.-Y., Robert A. 2009 Cellular encapsulation in the eastern subterranean termite, *Reticulitermes flavipes* (Isoptera), against infection by the entomopathogenic fungus *Metarhizium anisopliae*. *J Invertebr Pathol* **101**(3), 234-241. (doi:10.1016/j.jip.2009.05.008).
4. Dillon R.J., Charnley A.K. 1985 A technique for accelerating and synchronising germination of conidia of the entomopathogenic fungus *Metarhizium anisopliae*. *Arch Microbiol* **142**, 204-206. (doi:10.1007/BF00447069).
5. Barreto L.P., Luz C., Mascarin G.M., Roberts D.W., Arruda W., Fernandes E.K. 2016 Effect of heat stress and oil formulation on conidial germination of *Metarhizium anisopliae* s.s. on tick cuticle and artificial medium. *J Invertebr Pathol* **138**, 94-103. (doi:10.1016/j.jip.2016.06.007).
6. Kramm K.R., West D.F. 1982 Termite pathogens: Effects of ingested *Metarhizium*, *Beauveria*, and *Gliocladium* conidia on worker termites (*Reticulitermes* sp.). *J of Invertebr Pathol* **40**(1), 7-11. (doi:10.1016/0022-2011(82)90030-1).

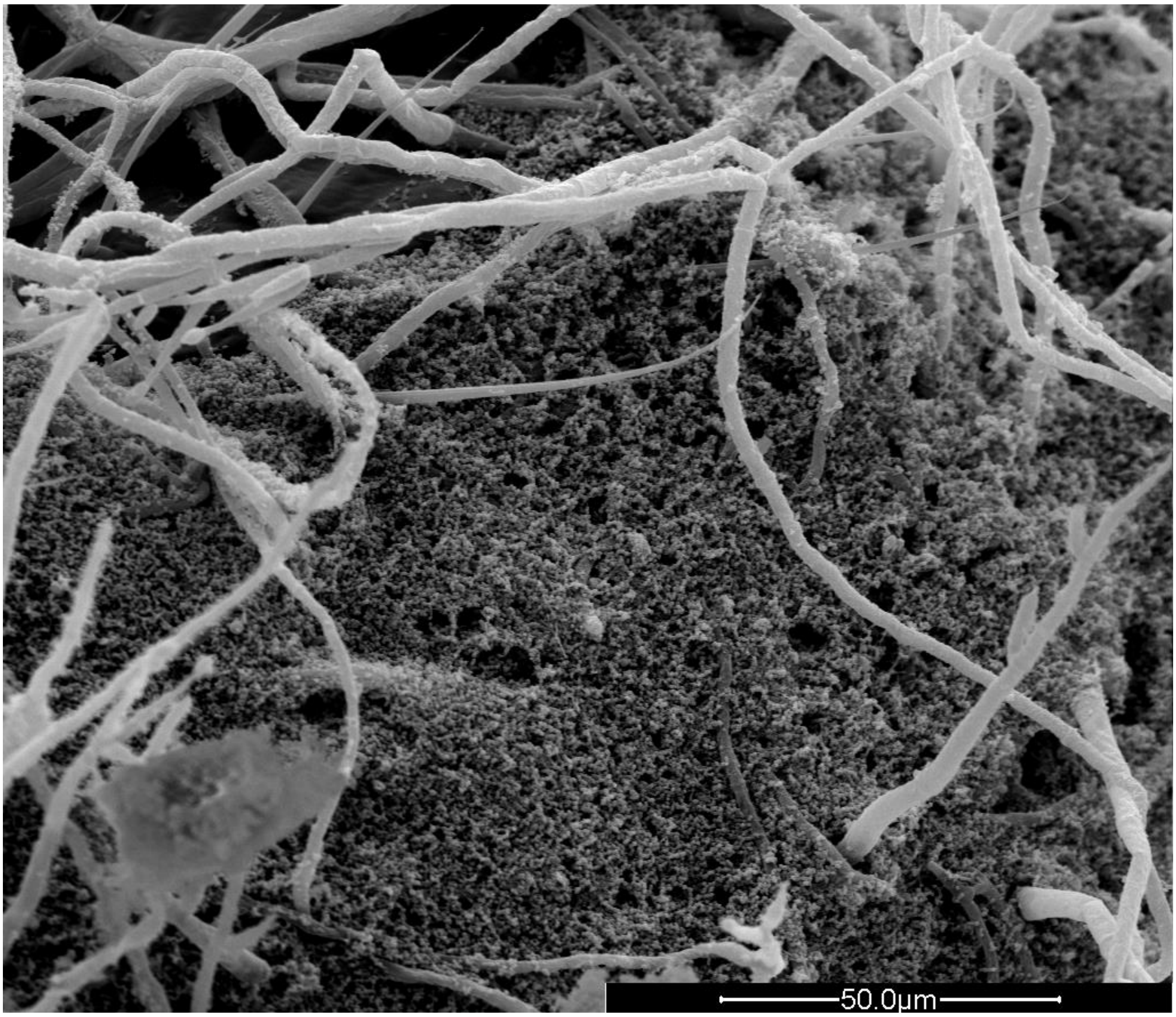


**Figure S2. Dense fungal and bacterial growth on the cuticle of a *R. flavipes* worker at 72 hours after exposure (approximately 2 days after death). 1500x.**

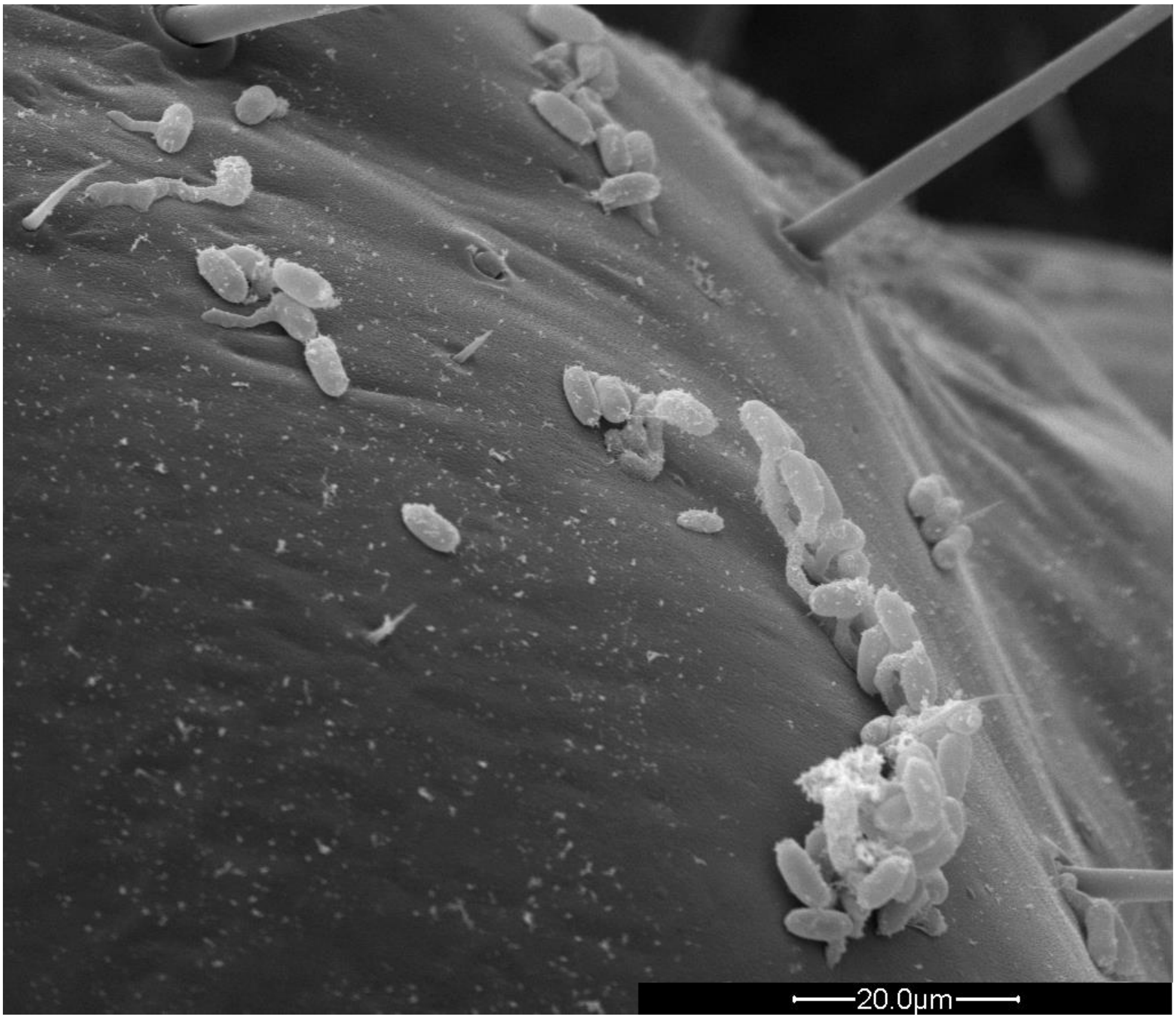


**Figure S3. Conidial germination on the cuticle of a *R. flavipes* worker at 10 hours after exposure. 2500x.**

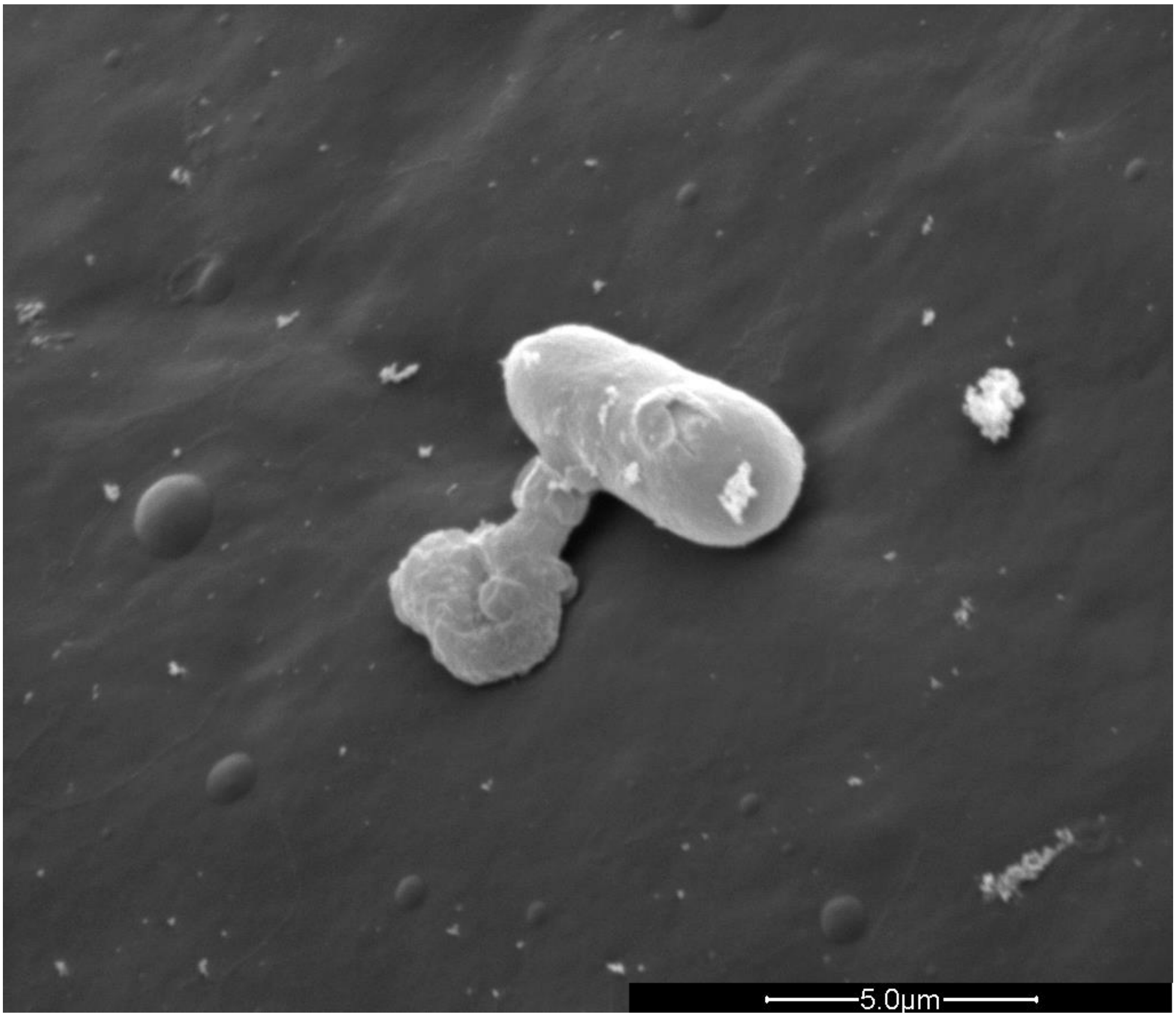


**Figure S4. Conidium with a germ tube and appressorium 12 hours after exposure. 12000x.**

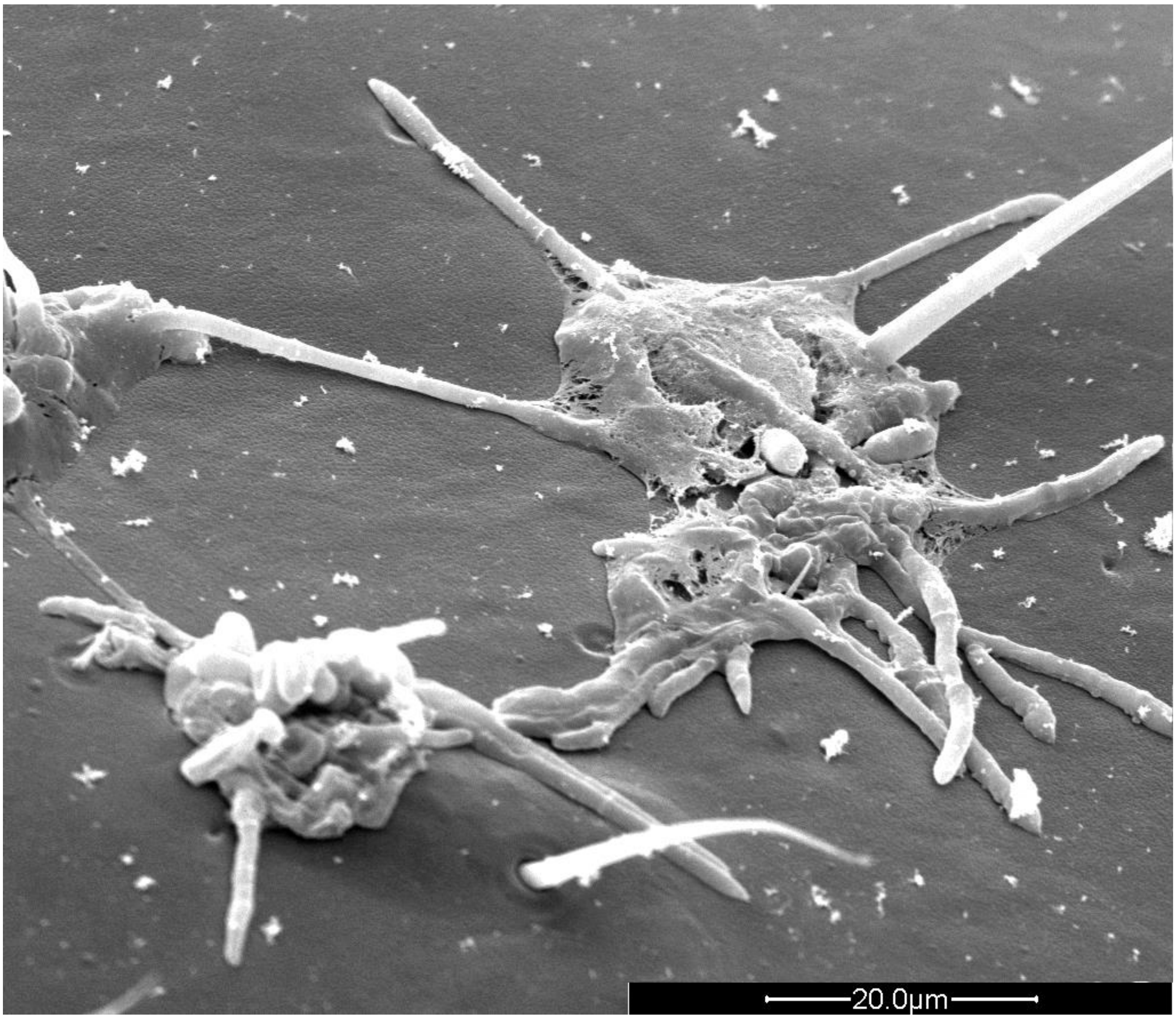


**Figure S5. Fungal growth on a *R. flavipes* worker cuticle 48 hours after exposure (death occurred between 24 and 48 hours after exposure). 300x.**

**Table S1. z and p values from post hoc pairwise comparisons (using Tukey correction) of the amount of grooming in all 8 treatments. Statistically significant differences are indicated in bold.**

| | | *M.a+* | | | | *M.a-* | | | |
|---|---|---|---|---|---|---|---|---|---|
| | | 2 h | 12 h | 15 h | 20 h | 2 h | 12 h | 15 h | 20 h |
| *M.a+* | 2 h | | | | | **z = 4.844<br>p < 0.001** | **z = 3.357<br>p = 0.01733** | **z = 4.504<br>p < 0.001** | **z = 3.765<br>p = 0.00384** |
| | 12 h | **z = 5.533<br>p < 0.001** | | | | **z = 9.126<br>p < 0.001** | **z = 7.834<br>p < 0.001** | **z = 8.828<br>p < 0.001** | **z = 8.056<br>p < 0.001** |
| | 15 h | z = -0.063<br>p = 1.00000 | **z = -5.511<br>p < 0.001** | | | **z = 4.752<br>p < 0.001** | **z = 3.273<br>p = 0.02266** | **z = 4.417<br>p < 0.001** | **z = 3.686<br>p = 0.00531** |
| | 20 h | z = -2.064<br>p = 0.42916 | **z = -7.717<br>p < 0.001** | z = -1.970<br>p = 0.49205 | | **z = 3.293<br>p = 0.02129** | z = 1.722<br>p = 0.66382 | z = 2.936<br>p = 0.06320 | z = 2.202<br>p = 0.34119 |
| *M.a-* | 2 h | | | | | | | | |
| | 12 h | | | | | z = 1.427<br>p = 0.83935 | | | |
| | 15 h | | | | | z = 0.327<br>p = 0.99998 | z = -1.100<br>p = 0.95503 | | |
| | 20 h | | | | | z = 0.923<br>p = 0.98318 | z = -0.477<br>p = 0.99974 | z = 0.602<br>p = 0.99880 | |

**Table S2. z and p values from post hoc pairwise comparisons (using Tukey correction) of the grooming intensity in all 8 treatments. Statistically significant differences are indicated in bold.**

| | | *M.a+* | | | | *M.a-* | | | |
|---|---|---|---|---|---|---|---|---|---|
| | | 2 h | 12 h | 15 h | 20 h | 2 h | 12 h | 15 h | 20 h |
| *M.a+* | 2 h | | | | | z = 1.990<br>p = 0.45148 | z = 1.766<br>p = 0.60864 | z = 1.567<br>p = 0.74189 | z = 2.117<br>p = 0.36841 |
| | 12 h | **z = 3.299<br>p = 0.01834** | | | | **z = 3.685<br>p = 0.00487** | **z = 3.839<br>p = 0.00269** | **z = 3.391<br>p = 0.01328** | **z = 3.950<br>p = 0.00169** |
| | 15 h | **z = 7.915<br>p < 0.001** | **z = 5.757<br>p < 0.001** | | | **z = 6.147<br>p < 0.001** | **z = 6.801<br>p < 0.001** | **z = 6.047<br>p < 0.001** | **z = 6.592<br>p < 0.001** |
| | 20 h | **z = 8.077<br>p < 0.001** | **z = 5.873<br>p < 0.001** | z = 0.084<br>p = 1.00000 | | **z = 6.198<br>p < 0.001** | **z = 6.860<br>p < 0.001** | **z = 6.107<br>p < 0.001** | **z = 6.633<br>p < 0.001** |
| *M.a-* | 2 h | | | | | | | | |
| | 12 h | | | | | z = 0.438<br>p = 0.99983 | | | |
| | 15 h | | | | | z = 0.424<br>p = 0.99986 | z = 0.012<br>p = 1.00000 | | |
| | 20 h | | | | | z = 0.010<br>p = 1.00000 | z = -0.449<br>p = 0.99979 | z = -0.432<br>p = 0.99984 | |

**Table S3. z and p values from post hoc pairwise comparisons (using Tukey correction) of the probability of not being cannibalised. Statistically significant differences are indicated in bold.**

| | | *M.a+* | | | | *M.a-* | | | |
|---|---|---|---|---|---|---|---|---|---|
| | | 2 h | 12 h | 15 h | 20 h | 2 h | 12 h | 15 h | 20 h |
| *M.a+* | 2 h | | | | | z = -0.442<br>p = 0.99980 | z = -0.442<br>p = 0.99980 | z = -0.442<br>p = 0.99980 | z = -0.510<br>p = 0.99949 |
| | 12 h | z = 2.355<br>p = 0.22934 | | | | z = 1.760<br>p = 0.60400 | z = 1.760<br>p = 0.60383 | z = 1.760<br>p = 0.60377 | z = 1.669<br>p = 0.66745 |
| | 15 h | **z = 4.550<br>p < 0.001** | **z = 5.233<br>p < 0.001** | | | **z = 3.958<br>p = 0.00153** | **z = 3.958<br>p = 0.00171** | **z = 3.958<br>p = 0.00167** | **z = 3.867<br>p = 0.00262** |
| | 20 h | **z = 5.469<br>p < 0.001** | **z = 6.745<br>p < 0.001** | **z = 3.006<br>p = 0.04437** | | **z = 4.896<br>p < 0.001** | **z = 4.896<br>p < 0.001** | **z = 4.896<br>p < 0.001** | **z = 4.809<br>p < 0.001** |
| *M.a-* | 2 h | | | | | | | | |
| | 12 h | | | | | z = 0.000<br>p = 1.00000 | | | |
| | 15 h | | | | | z = 0.000<br>p = 1.00000 | z = 0.000<br>p = 1.00000 | | |
| | 20 h | | | | | z = 0.068<br>p = 1.00000 | z = 0.068<br>p = 1.00000 | z = 0.068<br>p = 1.00000 | |
